# Host-Adapted Strains of *Spodoptera frugiperda* Hold and Share a Core Microbial Community Across the Western Hemisphere

**DOI:** 10.1101/2021.12.03.471132

**Authors:** Nathalia C. Oliveira, Pedro A.P. Rodrigues, Fernando L. Cônsoli

**Author notes:** Nathalia Cavichiolli de Oliveira –; Pedro Augusto da Pos Rodrigues –. **Correspondence** Fernando L. Cônsoli, Insect Interactions Laboratory, Department of Entomology and Acarology, Luiz de Queiroz College of Agriculture, University of São Paulo, Piracicaba, São Paulo, Brazil.

## Abstract

The fall armyworm *Spodoptera frugiperda* is an important polyphagous agricultural pest in the Western Hemisphere and currently invasive to countries of the Eastern Hemisphere. This species has two host-adapted strains named “rice” and “corn” strains. Our goal was to identify the occurrence of core members in the gut bacterial community of Fall armyworm larvae from distinct geographical distribution and/or host strain. We used next-generation sequencing to identify the microbial communities of *S. frugiperda* from corn fields in Brazil, Colombia, Mexico, Panama, Paraguay, and Peru, and rice fields from Panama. The larval gut microbiota of *S. frugiperda* larvae did not differ between the host strains neither was it affected by the geographical distribution of the populations investigated. Our findings provide additional support for *Enterococcus* and *Pseudomonas* as core members of the bacterial community associated with the larval gut of *S. frugiperda*, regardless of the site of collection or strain, suggesting that these bacteria may maintain true symbiotic relationships with the fall armyworm. Further investigations are required for a deeper understanding of the nature of this relationship.

## Introduction

The complexity and wide variety of host-microbe interactions are increasingly evident through new molecular techniques and the improvement of bioinformatic analysis tools. The advancement of understanding of this topic has brought support to some hypotheses, and challenged others. An example is the discussion on whether the gut microbiota is relevant for all animals [1]. The gut is a rich environment for holding a variety of host – microorganism associations, and the gut microbiota has been shown to play crucial roles in a wide range of aspects of host physiology, morphology and ecology. The insect gut microbiota can influence intra and interspecific interactions, such as sexual behavior [2, 3] and the relationship between host plants and natural enemies [4]. It also plays a key role in insect adaptation to their environment by providing essential nutrients [5, 6] and/or boosting the host immune response to parasites and pathogens [7, 8]. In addition, microbial symbionts can contribute to hosts by detoxifying xenobiotics as insecticides [9–12].

Such range of beneficial contributions has led to the establishment of true mutualistic associations in several groups of hemipterans, dipterans, blattids, and coleopterans, among others [9, 13–17]. Lepidopteran larvae, however, have been thought not to have established mutualistic associations with their gut-associated bacteria. Some studies demonstrated the survival, development time, and weight gain were not affected in antibiotic-fed larvae [18]. Additionally, the lack of special regions in the gut to house microorganisms has been argued as a strong limitation for the establishment of true associations with free-living microbes [19]. The harshness of the extremely alkaline conditions of the gut to most microorganisms also represents an unfavorable condition for establishing microbial associations [20]. Finally, the high variation in the composition of the microbial community driven by host plants would difficult the occurrence of associations that could hold through the required evolutionary time in order to allow the selection and establishment of true gut residents [21]. Nevertheless, other studies have shown that even in hostile environments as the midgut of lepidopteran larvae, there are evidence of gut colonization by certain bacterial groups [22–24]. In addition, gut-resident bacteria of lepidopteran larvae were demonstrated to play important physiological roles for their hosts [25, 26]; besides, the continuous association with their hosts for some of these microbes has been proved as they are horizontally transmitted [27].

Controversial topics in the scientific literature are always an invitation to new studies aiming at better understanding and clarification of the topic. The debated existence of true gut-associates in lepidoptera is a subject that needs further clarification due to two important contexts it is placed in. First, its remarkable relevance to the understanding of how microbial associations can influence host phenotypes [28], and insects have provided simple models for the clarification of fundamental principles in host-microbe interactions [29, 30], with a great potential to assist in unravelling complex systems such as in mammalians. Second, lepidopterans are yet the major group of agricultural pests, causing severe losses in food production, posing serious threats to food security [31–33]. Understanding the diversity and function of gut - microbes associations can lead to the development of new strategies for herbivore control.

In the present study we have chosen a lepidopteran species that is important both in the ecological and in the economic context to investigate the existence of true gut associates of lepidopteran larvae. *Spodoptera frugiperda* is an important agricultural pest in the Western Hemisphere and is currently invasive to countries in Africa, Asia, and Oceania [34–38]. *Spodoptera frugiperda* is highly polyphagous, feeding on more than 300 host plants [39]. This species is actually a complex composed of two distinct strains known as the rice (RS) and corn (CS) strains. The two strains are morphologically identical, with clear differences in host preference, susceptibility to insecticides and transgenic crops (*Bacillus thuringiensis*), composition of sex pheromone and mating behavior [40–47]. Genomic analysis of the host-adapted strains of *S. frugiperda* identified several genes involved in the chemodetection of non-volatile molecules and detoxification of xenobiotics showing signatures of positive selection, suggesting their contribution to *S. frugiperda* host plant preferences [48]. Some of these genomic variations between host strains of *S. frugiperda* were also detected at the transcriptional level, including those involved in xenobiotic metabolism [49].

Genetic studies suggest that population structure of *S. frugiperda* in the Western Hemisphere shows more variation within *S. frugiperda* populations than between populations of different locations, indicating a significant gene flow [50, 51]. The Mexican populations, on the other hand, have proven to be the most different, suggesting limited migratory interactions with foreign populations [52, 53]. The population genetic structure of Brazilian populations of *S. frugiperda* is partially based on host plants, with rice populations, which are basically represent by rice strain individuals, having a strong effect on the overall genetic structure of fall armyworm populations in Brazil [54].

Therefore, in this study we aim to verify the existence of bacterial groups that remain associated with the gut microbial community of *S. frugiperda* larvae regardless of the geographical region or host plant used. So, we sampled and sequenced the gut microbiota of fall armyworm larvae from corn and rice fields across the American continent. Larvae were genotyped as rice or corn strain, and the structure of the bacterial gut community was checked based on the geographical origin of the larvae, host-adapted strain and/or host plant used. Despite the variation expected due to uncontrolled and unforeseen environmental factors, the field conditions may provide essential information on potential symbionts that could be ecologically important to their hosts in their natural habitats.

## Material and Methods

### Sampling and strains identification

Larvae of *Spodoptera frugiperda* with 2.5-3.0 cm in length were collected from corn and/or rice fields during 2016-17 in Brazil (13.8224° S, 56.0835° W), Colombia (4.5709° N, 74.2973° W), Mexico (23.6345° N, 102.5528° W), Panama (8.5380° N, 80.7821° W), Paraguay (23.4425° S, 58.4438° W), and Peru (9.1900° S, 75.0152° W), and stored in absolute ethanol. Once in the laboratory, larvae had the width of the head capsule measured, and only those larvae with head capsule width within the limits of size of 5^th^ and 6^th^ instars [55] were further dissected for gut collection. Dissections were carried after surface sterilization under aseptic conditions in a laminar flow hood. The larval digestive tract was carefully removed, washed in sterile saline and further used in metabarcoding analysis of the gut microbiota. The remaining carcass was used for host strain identification.

*Spodoptera frugiperda* were genotyped for strain identification using the mitochondrial *cytochrome oxidase I* (*COI*) gene as a marker. DNA was extracted using the genomic DNA preparation protocol from RNALater™, with modifications. The carcass obtained from dissected larvae was placed in 2 mL tubes with 750 μL digestion buffer (60 mM Tris pH 8.0, 100 mM EDTA, 0.5% SDS) and proteinase K (500 μg/mL), macerated using pestle, and mixed well by inversion. Samples were incubated overnight at 55°C. Afterwards, 750 μL of phenol:chloroform (1:1) was added and rapidly inverted for 2 min. Samples were centrifuged at high speed for 10 min. The aqueous layer was collected and phenol:chloroform extraction was repeated twice before a final extraction with chloroform. The aqueous layer was collected, added to 0.1 volume of 3M sodium acetate (pH 5.2) and an equal volume of 95% ethanol. Samples were then mixed by inversion, incubated for 40 min at −80°C before centrifugation (27,238 *g* x 30 min x 4°C). The pellet obtained was washed twice with 1 mL of 85% ice-cold ethanol, centrifuged for 10 min after each wash, and dried at 60°C during 5-10 min in a SpeedVac. Finally, the pellet was resuspended in nuclease-free water. DNA concentration and quality were estimated by spectrophotometry and standard DNA agarose gel electrophoresis [56].

Polymerase chain reactions (PCR) for partial amplification of the mitochondrial *COI* gene was conducted using the primer set JM76 (5’-GAGCTGAATTAGGRACTCCAGG-3’) and JM77 (5’-ATCACCTCCWCCTGCAGGATC-3’), to produce an expected amplicon of 569 base pairs (bp) [57]. The PCR mixture contained 100-150 ng of gDNA, 1.5 mM of MgCl_2_, 1x PCR buffer, 0.2 mM of each dNTP, 0.32 μM of each primer and 0.5U of GoTaq^®^ DNA Polymerase (Promega) in a total volume of 25 μL. The thermocycling condition was 94°C x 1 min (1x), followed by 33 cycles at 92°C x 45 s, 56°C x 45 s, and 72°C x 1 min, and one cycle at 72°C x 3 min for final extension. Amplicons were then subjected to restriction analysis using the *Msp*I (HpaII) restriction endonuclease. Samples were gently mixed, centrifuged for a few seconds and incubated overnight at 37°C. Subsequently, digestion and the resulting products were verified using a 1.5% agarose gel electrophoresis. The corn strain (*CS*) was identified from restriction analyses yielding two fragments (497bp and 72bp), while restriction analyses that produced no digestion identified the rice strain (*RS*) [57].

### DNA extraction, amplification and 16S rDNA sequencing

The midgut obtained from dissected larvae were individually powdered in liquid nitrogen, and genomic DNA was extracted using the Wizard Genomic DNA Purification Kit (Promega), following the manufacturer’s recommendations. The quality, integrity and purity of the DNA obtained was measured by spectrophotometry and agarose gel electrophoresis as before. DNA samples were stored in −20°C and sent for library construction, normalization and sequencing in the Center for Functional Genomics (http://www.esalq.usp.br/genomicafuncional/), one of the multiusers laboratories of our institution. Paired-end reads were generated after amplifying the v3-v4 region of 16S rRNA gene (approximately 550 bp) using the Nextera XT DNA Library Preparation Kit (Illumina) for paired-end (2x 300 bp) sequencing in the Illumina MiSeq platform.

### Sequences analyses

Illumina adapters at the 3’ end of the reads were removed using *Cutadapt* [58]. The bioinformatics analyses of the gut microbiome were performed with QIIME2 v. 2020.2.0 [59]. Raw sequence data were quality filtered with *q2-dada2* plugin for filtering phiX reads and chimeric sequences [60]. In order to remove low quality regions from quality filter reads, *dada2 denoise-single* method trimmed off the first 18 nucleotides of the forward reads and 22 nucleotides from the reverse reads. It also truncated each sequence at position 290 in the forward and 220 in the reversed reads. These positions were chosen based on visual inspection of plotted quality scores from demultiplexed reads. A phylogeny was estimated with *SEPP* [61] as implemented in the *q2-fragment-insertion* QIIME2 plugin. All amplicon sequence variants (ASVs) were aligned with *feature-classifier classify-sklearn* against the SILVA-132-99 database [62] that was trained with a Naïve Bayes classifier [63] on the Illumina 16S rRNA gene primers targeting the V3–V4 region.

The downstream analysis was performed in the MicrobiomeAnalyst web platform (https://www.microbiomeanalyst.ca/) [64] and in R (version 4.0.4) [65]. Data were filtered keeping ASV with minimum count of four (4) per library and low count filter based on 20% prevalence across samples. Data were rarefied to the minimum library size (1155 reads), before any statistical comparisons. Rarefaction curves were based on the relationship between number of ASVs and number of sequences. Alpha diversity analysis was measured by the observed species and Shannon index. The results were plotted across samples and showed as box plots for each group. Beta diversity was investigated through principal components analysis (PCoA) using unweighted and weighted UniFrac distances, and through hierarchical clustering analysis using unweighted UniFrac distances.

We used PERMANOVA to test the strength and statistical significance of sample groupings based on generalized weighted UniFrac distances. This distance contains an extra parameter α (set at α=0.5) to control the weight of abundant lineages, so the distance is not dominated by highly abundant lineages. When differences were found between samples distances, a *post-hoc* analysis was performed with the package *pairwise.adonis* to identify differences among treatments and verify the adjusted *p* value [66]. As PERMANOVA assumes homogeneity of variances, we used *betadisper*, a multivariate analogue of Levene’s test, as implemented in R to verify whether differences between groups in terms of their centroids are not due to differences in variances. Analysis of similarity (ANOSIM) was used when there was heterogeneity of variance among groups. In our sample set we had basically 3 groups: (i) countries that presented both strains in corn plants, (ii) countries with only the corn strain in corn plants and (iii) Panama with both strains in corn plants and only the rice strain in the rice plant. Since our design is unbalanced, we performed separate analyses to properly grasp our data. First, we excluded the samples that had rice as host plant, thus only the variables “strain” and “country” were considered. To test the effect of country and host plant, we excluded the corn strain from the analysis, considering only the rice strain, and performed multilevel pairwise comparison using Adonis (PERMANOVA) from package *vegan* with adjusted *p*-values.

To visualize taxa abundance across the different groups, taxa plots were constructed based on phyla and genera. The core microbiome analysis was defined as the genera present in 50% or more of the samples and showing a relative abundance of 0.05% in each library. The differential abundance analysis was also analyzed using *DESeq2* methods [67]. Pattern Search was used to identify which features were correlated with the core microbiome in the gut microbial community. Pearson r was the distance measure used using the MicrobiomeAnalyst tool [64].

To cluster our samples groups into distinct ‘metacommunities’, we performed Dirichlet multinomial mixtures using the *get.communitytype* function [68] after exportation of biom ASV table from qiime2 to Mothur (v.1.44.3) and the selection of subsamples with *subsample*=*1000*, excluding low abundance samples that might be a result of artifact operational units and/or variation due to rare taxons (“singletons”). The best fitting number of metacommunities was obtained by selecting the minimum local Laplace value obtained after five iterations.

## Results

A total of 63 *S. frugiperda* individuals, 8 *RS* and 45 *CS* were used in our analyses. Except for 8 specimens from Panama that were collected on rice, all other samples were collected in corn fields. Out of the 63 specimens analyzed, 21 were from Brazil (*CS*=18; *RS*=3), nine from Colombia (*CS*=8; *RS*=1), eight from Mexico (*CS*=8), six from Paraguay (*CS*=3; *RS*=3), five from Peru (*CS*=3; *RS*=2), and 13 from Panama (6 from corn fields; *CS*=5, *RS*=1; and 8 from rice fields; *RS*=8).

Rarefaction analysis (Fig. S1) showed that sampling was adequate for an accurate characterization of the diversity and richness of the larval gut microbiota of *S. frugiperda*. Samples that failed to achieve adequate sampling depth were excluded from further analyses. There was no difference in alpha-diversity values between strains or among countries (Fig. 1) measure by observed species and Shannon diversity indices. The beta diversity measured by weighted Unifrac distances did not exhibit specific clustering based on the country of origin or *S. frugiperda* strains (Fig. 2).

**Fig. 1.**
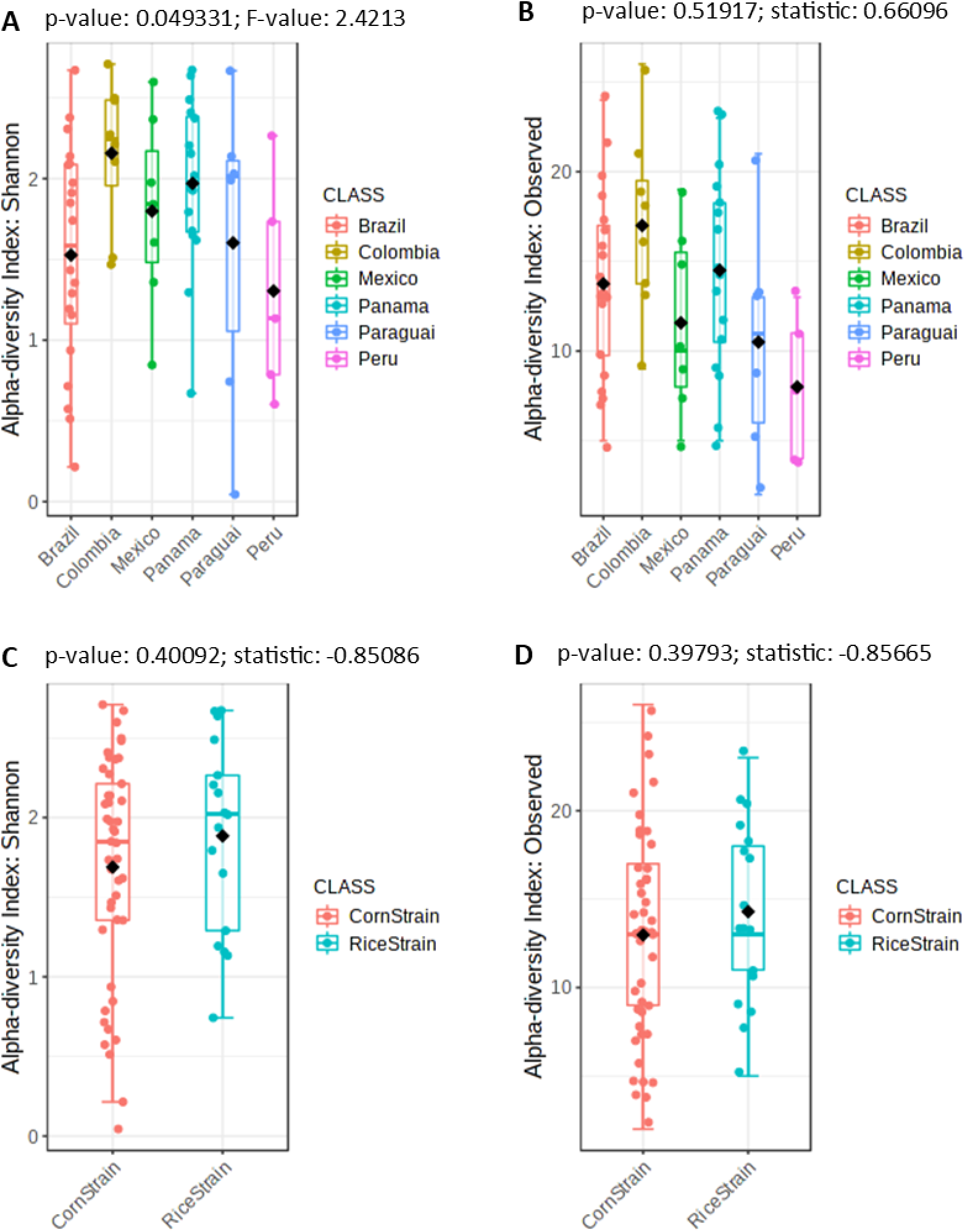
Alpha diversity index of Shannon index (A, C) and observed taxa (B, D) obtained for samples from the gut microbiota of the corn and rice strains of *Spodoptera frugiperda* larvae (C, D) from different countries (A, B). The statistical values from the Test t (pairwise comparison) and ANOVA (group comparison) are shown in which box.

**Fig. 2.**
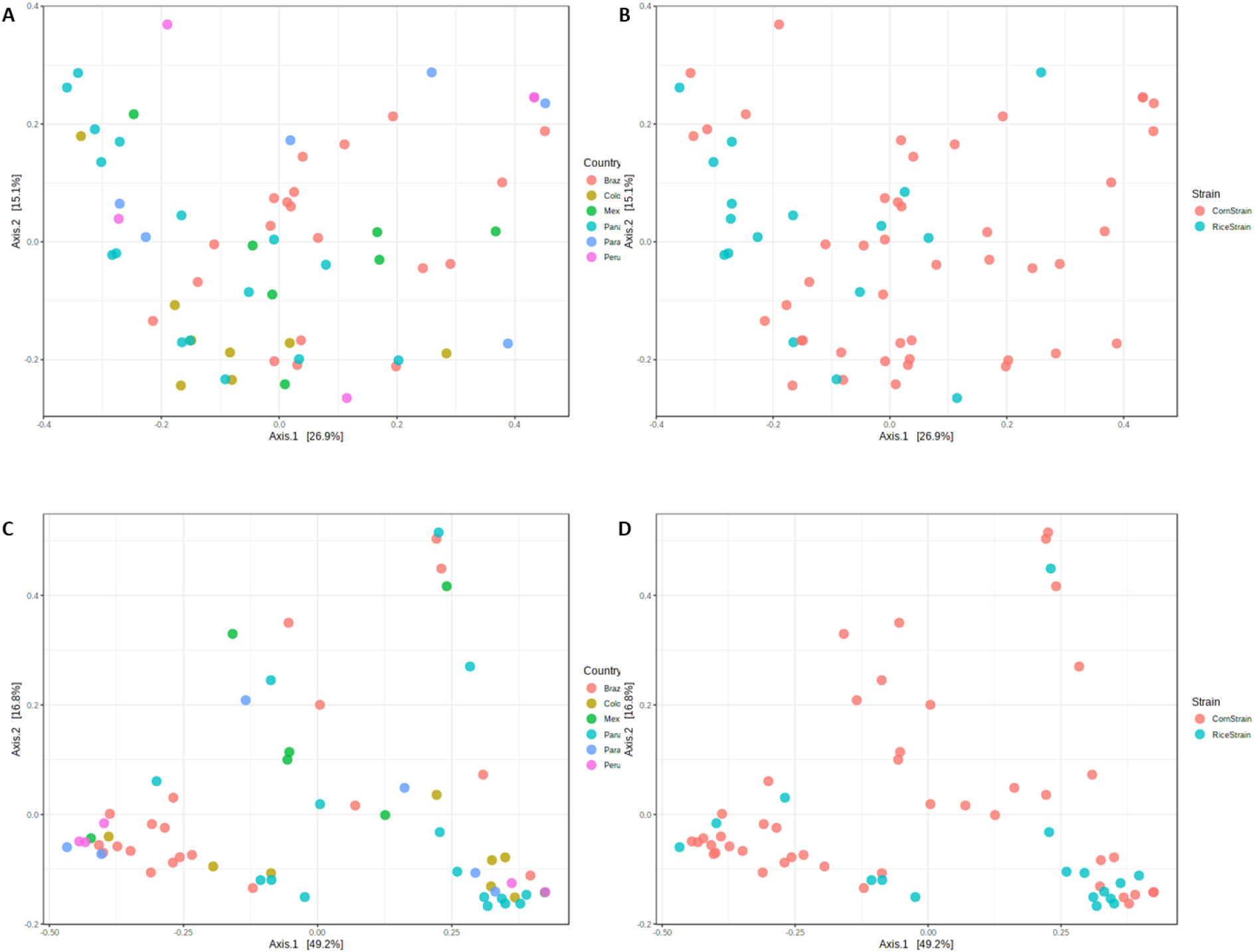
Principal coordinates analysis (PCoA) based on unweighted (A, B) and weighted (C, D) unifrac analysis of the midgut microbial community of the corn and rice strains of *Spodoptera frugiperda* larvae (B, D) from different countries (A, C). The statistical values from PERMANOVA are shown in each box.

When considering samples collected in maize, no differences in the composition of the gut microbial community between strains (*p*=0.215) (Table 1) nor among different countries considering the adjusted *p-* values (*p*-values < 0.05) were detected (Table 2). Betadisper showed that groups had the same dispersion, failing to reject the null hypothesis of homogeneous multivariate dispersions, meeting the assumption for Adonis (Table 1). It thus provided confidence to the PERMANOVA results, meaning the values obtained were not an artifact of heterogeneity of dispersions. Likewise, no differences were found between host plants (*p*=0.344) or country (*p*=0.0709) when considering only the rice strain (Table 3). Additionally, all replicates of metacommunity analyses resulted in the same pattern (K=1), meaning that according to the Dirichlet model there is not a clear pattern of grouping ASVs across samples.

**Table 1.**
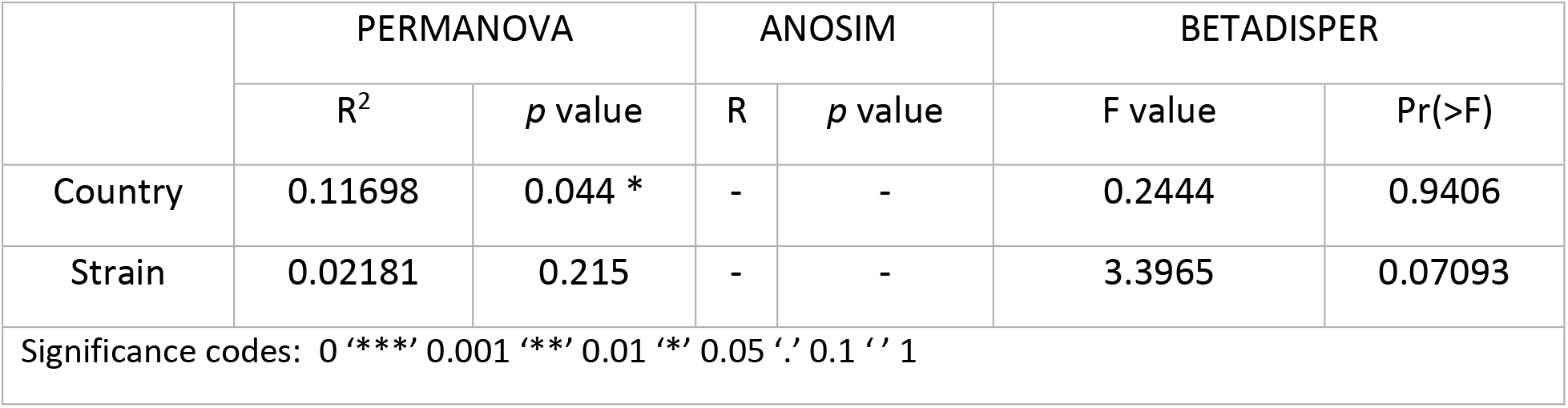
PERMANOVA and BETADISPER results from comparisons of the gut microbial communities among countries and *Spodoptera frugiperda* strains (corn and rice strains) excluding samples from rice plants using UniFrac (alpha 0.5) values.

**Table 2.** Post-hoc analysis of comparisons of the *Spodoptera frugiperda* gut microbial communities among countries using UniFrac (alpha 0.5) values.

| Pairs | Df | SumsOfSqs | F.Model | R2 | p.value | p.adjusted |
| --- | --- | --- | --- | --- | --- | --- |
| Colombia vs Brazil | 1 | 0.3806171 | 2.0309445 | 0.07245366 | 0.011 | 0.165 |
| Colombia vs Mexico | 1 | 0.2728639 | 1.3648260 | 0.08882795 | 0.080 | 1.000 |
| Colombia vs Panama | 1 | 0.2279787 | 1.1800870 | 0.07773915 | 0.196 | 1.000 |
| Colombia vs Paraguai | 1 | 0.3238154 | 1.6384779 | 0.12013642 | 0.095 | 1.000 |
| Colombia vs Peru | 1 | 0.3136700 | 1.6244547 | 0.12867524 | 0.060 | 0.900 |
| Brazil vs Mexico | 1 | 0.2544575 | 1.2752619 | 0.04675526 | 0.147 | 1.000 |
| Brazil vs Panama | 1 | 0.3235731 | 1.6516793 | 0.05973161 | 0.044 | 0.660 |
| Brazil vs Paraguai | 1 | 0.1472816 | 0.7425192 | 0.03000985 | 0.722 | 1.000 |
| Brazil vs Peru | 1 | 0.2553736 | 1.3015037 | 0.05355651 | 0.151 | 1.000 |
| Mexico vs Panama | 1 | 0.2217735 | 1.0281323 | 0.06841384 | 0.390 | 1.000 |
| Mexico vs Paraguai | 1 | 0.2483650 | 1.1092582 | 0.08461640 | 0.292 | 1.000 |
| Mexico vs Peru | 1 | 0.2227645 | 1.0045716 | 0.08368242 | 0.383 | 1.000 |
| Panama vs Paraguai | 1 | 0.2516586 | 1.1648641 | 0.08848281 | 0.254 | 1.000 |
| Panama vs Peru | 1 | 0.2287484 | 1.0730526 | 0.08887998 | 0.328 | 1.000 |
| Paraguai vs Peru | 1 | 0.1878847 | 0.8404890 | 0.08541130 | 0.553 | 1.000 |

**Table 3.** PERMANOVA, ANOSIM and BETADISPER results from comparisons of the gut microbial 600 communities of the *Spodoptera frugiperda* rice strain among countries and host plants using UniFrac (alpha 601 0.5) values.

|  | PERMANOVA |  | ANOSIM |  | BETADISPER |  |
| --- | --- | --- | --- | --- | --- | --- |
|  | R<sup>2</sup> | *p* value | R | *p* value | F value | Pr(>F) |
| Country | - | - | 0.266<br>4 | 0.07092<br>9 | 5.6096 | 0.00755 ** |
| Host Plant | 0.0614<br>9 | 0.344 | - | - | 2.1328 | 0.1635 |

Significance codes: 0 '\*\*\*' 0.001 '\*\*' 0.01 '\*' 0.05 '.' 0.1 ' ' 1

At the phylum level, the midgut of *S. frugiperda* was composed by *Proteobacteria*, *Firmicutes* and *Actinobacteria* (Fig. 3). There was no significant difference at the phylum level among countries or between strains. Taxa bar plots at the genus level indicated that individuals from the same country exhibited a high degree of variability in terms of bacteria taxa abundance (Fig. 4). *Klebsiella* and *Erysipelatoclostridium* were the taxa that differed among countries (Fig. 5), and the abundance of *Erysipelatoclostridium* also differed between *RS* and *CS* (Fig. 6).

**Fig. 3.**
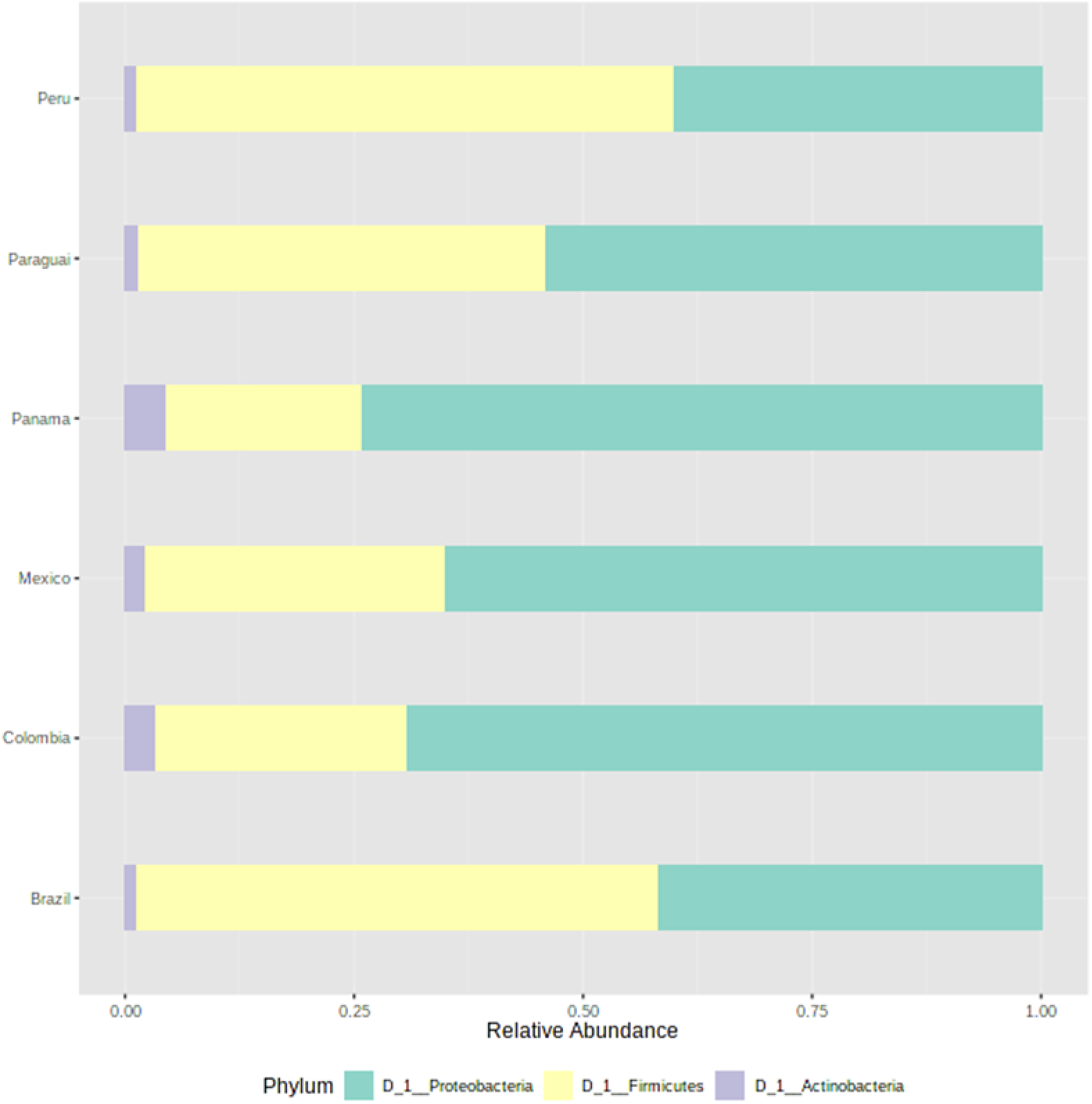
Taxonomic composition of the microbial community associated with the midgut of corn and rice strains of *Spodoptera frugiperda* larvae sampled in different countries at the phylum level.

**Fig. 4.**
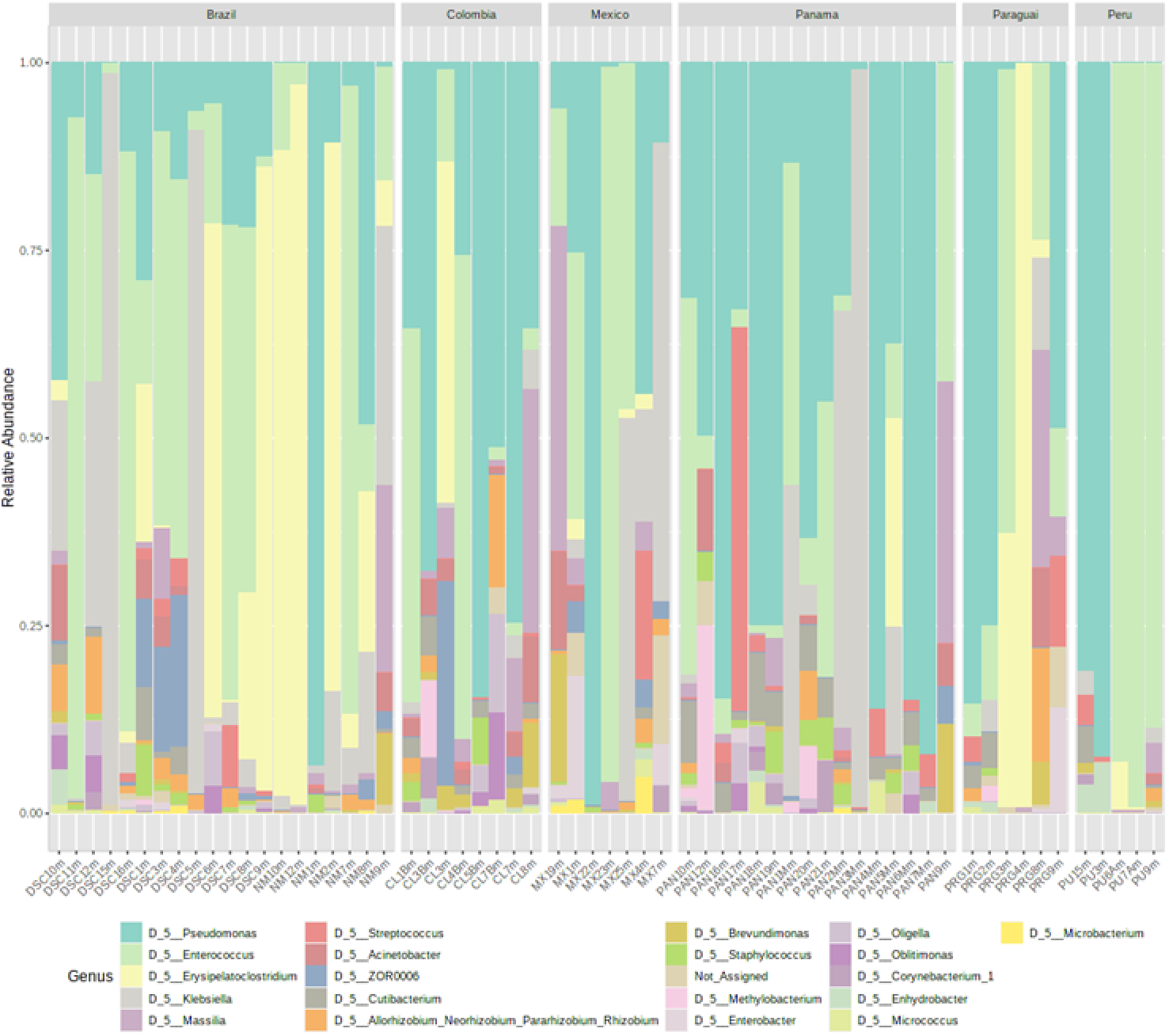
Taxonomic composition of the microbial community of the larval midgut of corn and rice strains of *Spodoptera frugiperda* at the genus level.

**Fig. 5.**
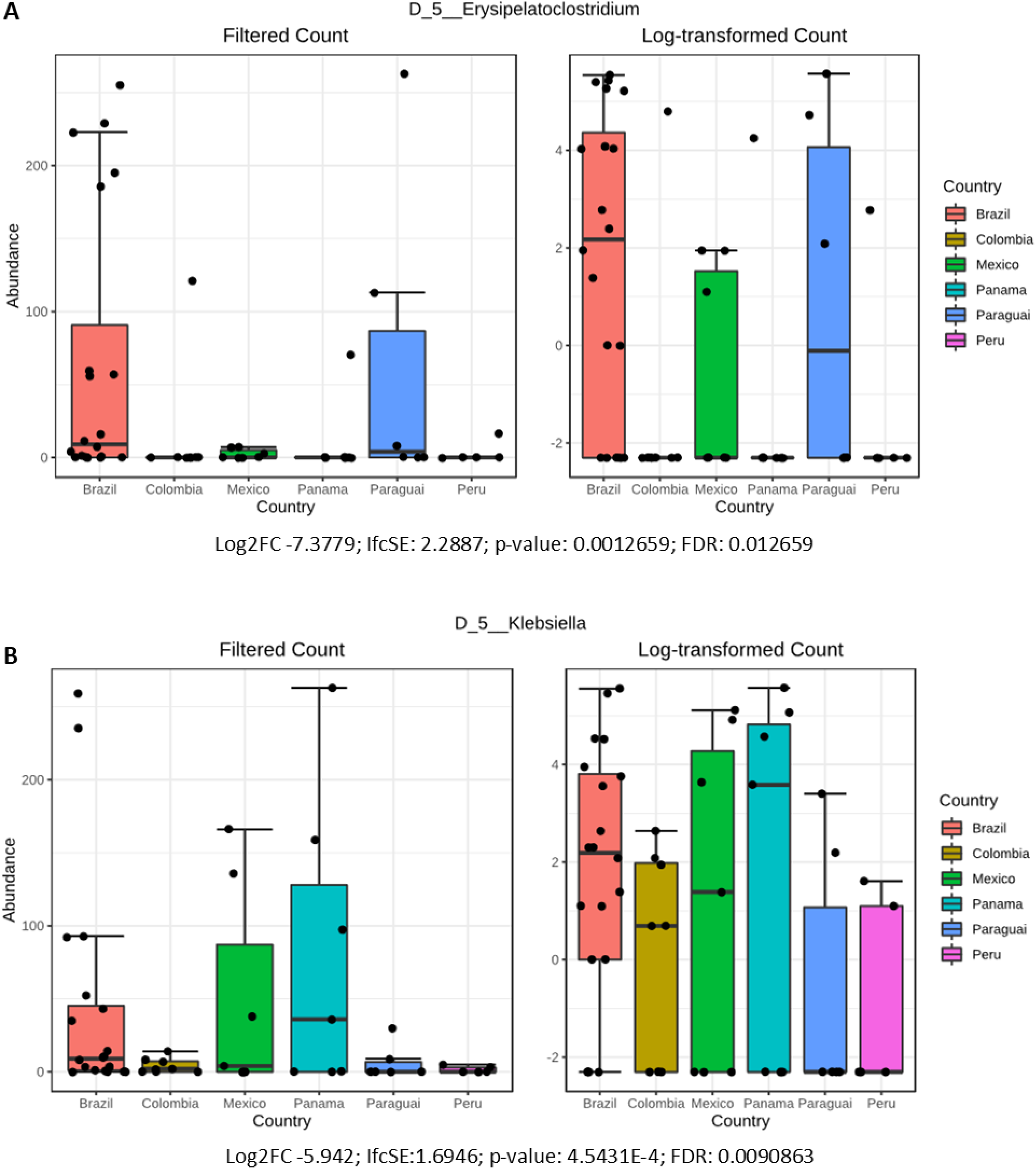
The abundance of *Klebsiella* and *Erysipelatoclostridium* as a differential feature of the microbiota associated with the larval midgut of *Spodoptera frugiperda* from different countries.

**Fig. 6.**
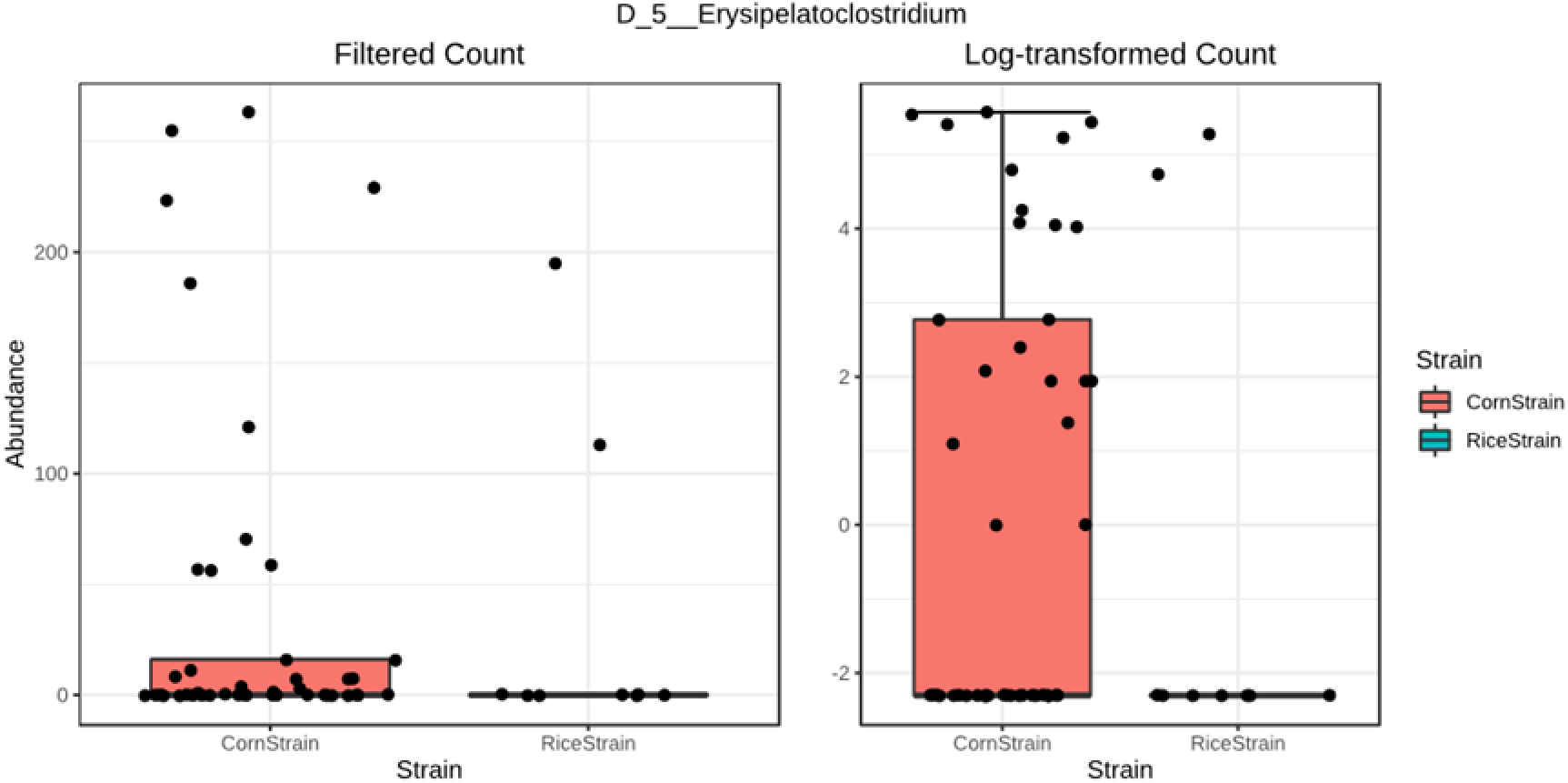
The abundance of *Erysipelatoclostridium* as a differential feature of the microbiota associated with the larval midgut of the corn and rice strains of *Spodoptera frugiperda*.

The bacterial core of the larval midgut of *S. frugiperda* at the genus level was composed by *Pseudomonas* and *Enterococcus*. Correlation analysis identified 10 genera that were positively correlated and 10 genera negatively correlated with *Pseudomonas*. However, only three genera were positively correlated, while 18 were negatively correlated with *Enterococcus* (Fig. 7).

**Fig. 7.**
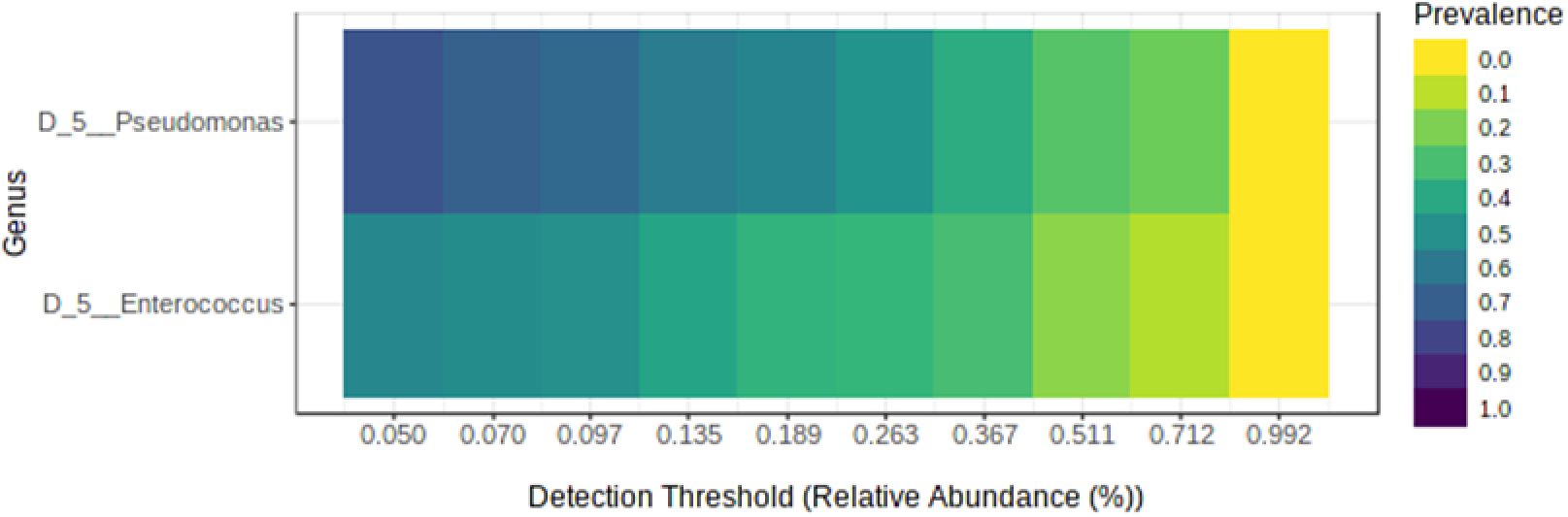
The core gut microbiota of *Spodoptera frugiperda* at the genus level identified by MicrobiomeAnalyst using the parameters sample prevalence (50 %) and relative abundance (0.5 %).

**Fig. 8.**
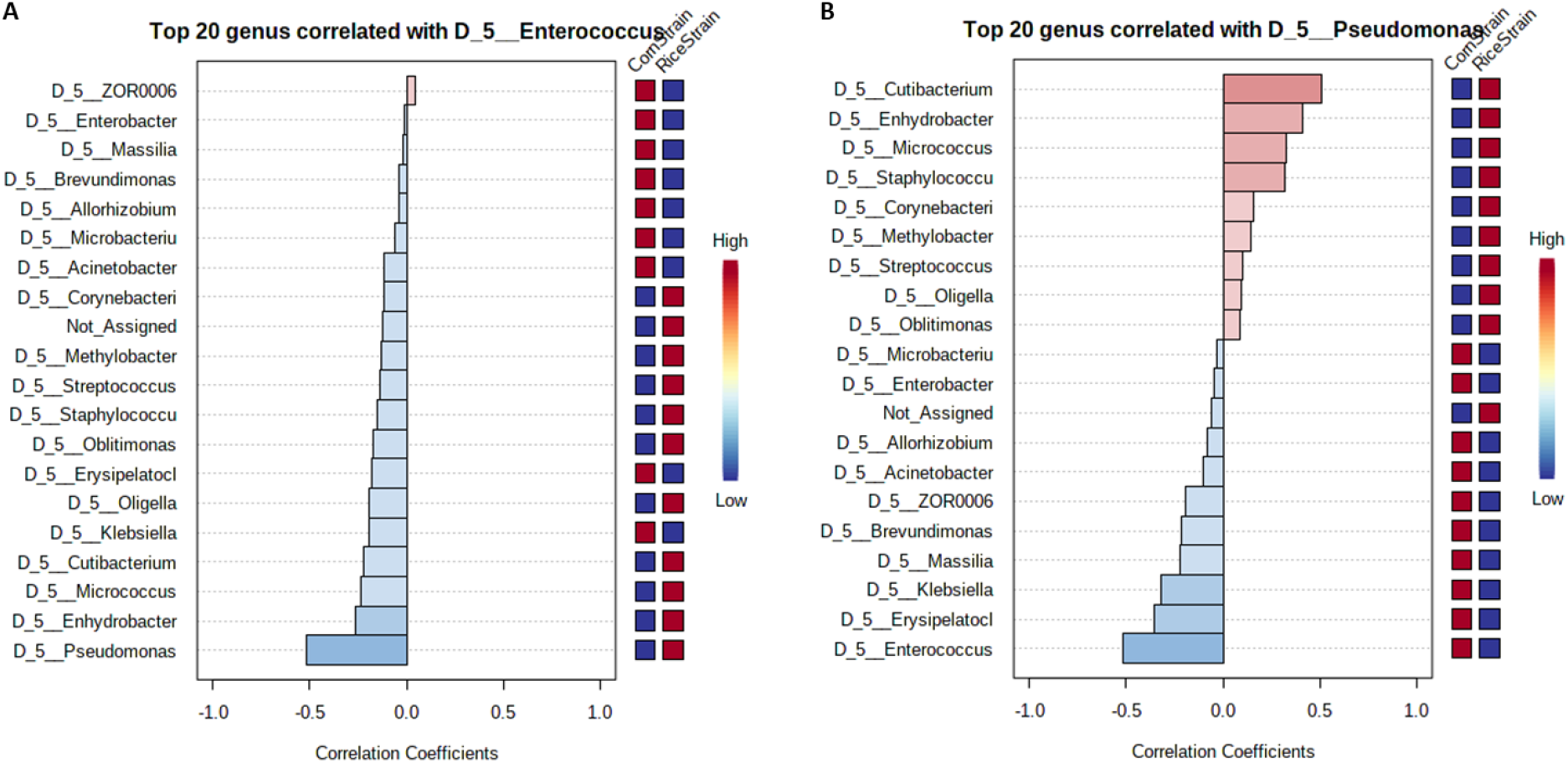
Pattern correlation analysis of the larval gut bacteria of *Spodoptera frugiperda* at the genus level. Red indicates positive correlation and blue indicates negative correlations with the presence of *Enterococcus* (A) or *Pseudomonas* (B).

## Discussion

Our results indicate that bacterial communities of the Fall armyworm larval midgut do not differ between strains collected from the same country nor among countries. These findings follow the pattern of the population genetic structure of *S. frugiperda* in the Western Hemisphere, where the majority of the genetic variability is within individual populations and not between populations, suggesting that populations of *S. frugiperda* functions as a panmictic population [50, 51].

As expected, we detected high variations in the composition of the gut microbiota among larvae. Such differences are likely to occur due to differences in corn varieties and associated endophytes, and soil type and associated microbiota, which also interact with plants and affect the plant endophyte community, ultimately interfering with the microbial composition of herbivores [69–71]. Variation in the microbiota from individual samples within treatments is commonly reported to several organisms, including species of Lepidoptera [72–74]. In humans, interindividual variation in the populations of gut microbes can be higher than 90% [75].

Obadia and collaborators [76] exploring the colonization of bacteria in the *Drosophila melanogaster* gut found that several strains of different species can maintain a stable association with the fly gut under laboratory conditions. They demonstrated that the establishment of bacteria in the gut works like a lottery and that stochastic factors generate alternative, stable states of gut colonization. Moreover, the resident species that have colonized the larval gut earlier, reduced the chances of subsequent colonization. Another interesting point raised concerning our study was that the peritrophic matrix in the midgut prevents the bacteria from attaching to the epithelial cells. Therefore, the lack of tissue attachment potentially makes these luminal populations less stable within the gut.

But regardless the high variation observed in the gut microbiota associated with the larval midgut of *S. frugiperda*, our analysis identified a core of bacteria despite the geographical origin of fall armyworm samples. The maintenance of a core independently of any interfering systemic effects points to the existence of bacterial associates with specific functions. In addition, the high variability in the composition of the midgut microbiota may allow for rapid host adaptation through rapid selection of microbiota suitable for contributing to the host under different stress conditions, such as abiotic factors, dietary resources, and risk of natural enemy attack [77].

The ASVs *Pseudomonas* and *Enterococcus* identified in this study as core members of the microbiota of the fall armyworm were also identified before as part of the core taxa associated with the gut of *S. frugiperda* larvae from corn fields [10, 77–80]. The high abundance of *Pseudomonas* in our samples suggests that this genus of bacteria could assist *S. frugiperda* larvae to overcome environmental stressors, particularly by aiding larvae to degrade natural and/or synthetic toxic xenobiotics. *Pseudomonas* capable to degrade several pesticides were recovered from the gut of laboratory-selected resistant lines (Almeida, Moraes, Trigo, Omoto, & Cônsoli, 2017), but also from field populations of *S. frugiperda* collected from several corn-producing areas in Brazil (Gomes, Omoto, & Cônsoli, 2020). *Pseudomonas* have also been demonstrated to degrade secondary metabolites in the gut of a coleopteran host [81]. Additionally, *Pseudomonas* abundance increased in the gut of *Plutella xylostella* resistant to prothiofos when compared to susceptible larvae, and was also shown to have antagonistic activity to several species of entomopathogenic fungi through siderophore production as demonstrated in culture plates [82].

It is noteworthy that *Enterococcus* is the most prevalent and abundant group identified in the gut microbiota of *Spodoptera* species [78, 79, 83], and also the most active in the gut of *S. frugiperda* [84]. Additionally, it has been demonstrated that *Enterococcus mundtii* is effective in colonizing and forming biofilm in the gut of *Spodoptera littoralis* [22, 27]. There is also evidence that *E. mundtii* can be inherited by *S. littoralis* through vertical transmission [23]. Some species of *Enterococcus* produce antimicrobial peptides with high level of inhibitory activity against potential bacterial competitors [27], which may explain its prevalence when compared to other phylotypes in *S. frugiperda* gut communities, but also the high negative correlation of *Enterococcus* with the other bacterial species of the gut microbiota community of *S. frugiperda* in this study.

Overall, this study provided an extended view of the fall armyworm gut microbiota and supported the hypothesis that bacterial taxonomic compositions across different localities in the Western Hemisphere are similar to each other, presenting high inter-individual variance, and that there are no significant differences in gut microbiota composition between the host-adapted strains of *S. frugiperda*. Nevertheless, our findings provide further evidence that *Pseudomonas* and *Enterococcus* are true symbionts of *S. frugiperda* as they were identified in the gut microbiota of *S. frugiperda* larvae regardless the host plant and site of collection. Further investigations on the functional contribution of these species as members of the gut bacterial community of fall armyworm larvae is required for a deeper understanding of the nature of this relationship.

## Declarations

### Availability of data and material

Upon paper acceptance, the data will be archived and the data regarding the deposited database and information such as access numbers will be provided for all manuscript data.

### Code availability

Not applicable

### Contributions

N.C.O. and F.L.C. conceived the study and designed the experiment. N.C.O. processed the samples. N.C.O. and P.A.P.R. conducted the bioinformatics. FLC secured funds for the project. N.C.O. wrote the first draft of the manuscript. P.A.P.R. and F.L.C. revised and edited the initial draft. All authors approved the final version for publication.

### Additional declarations for articles in life science journals that report the results of studies involving humans and/or animals

Not applicable

### Ethics approval

Not applicable

### Consent to participate

All authors agree with the participation in this manuscript.

### Consent for publication

All authors agree with the manuscript submission to Microbial Ecology Journal.

## Acknowledgements

We are grateful to the São Paulo Research Foundation (FAPESP) (process 2011/50877-0) and the Ministry of Science, Technology and Innovation (Conselho Nacional de Desenvolvimento Científico e Tecnológico – CNPq: process 462140-2014/8) for the grant provided to the senior author. The HPC resources made available by the Superintendence of Information Technology of the University of São Paulo. We also thank FAPESP for the PhD student fellowship (2017/24377-7) provided to the first author. We also would like to thank our collaborators from Panama, Peru, Colombia, Paraguay and Brazil who helped us with the field collection of larval samples. This manuscript is one of the chapters of the PhD Thesis of the first author.

---

**Fig. S1.**
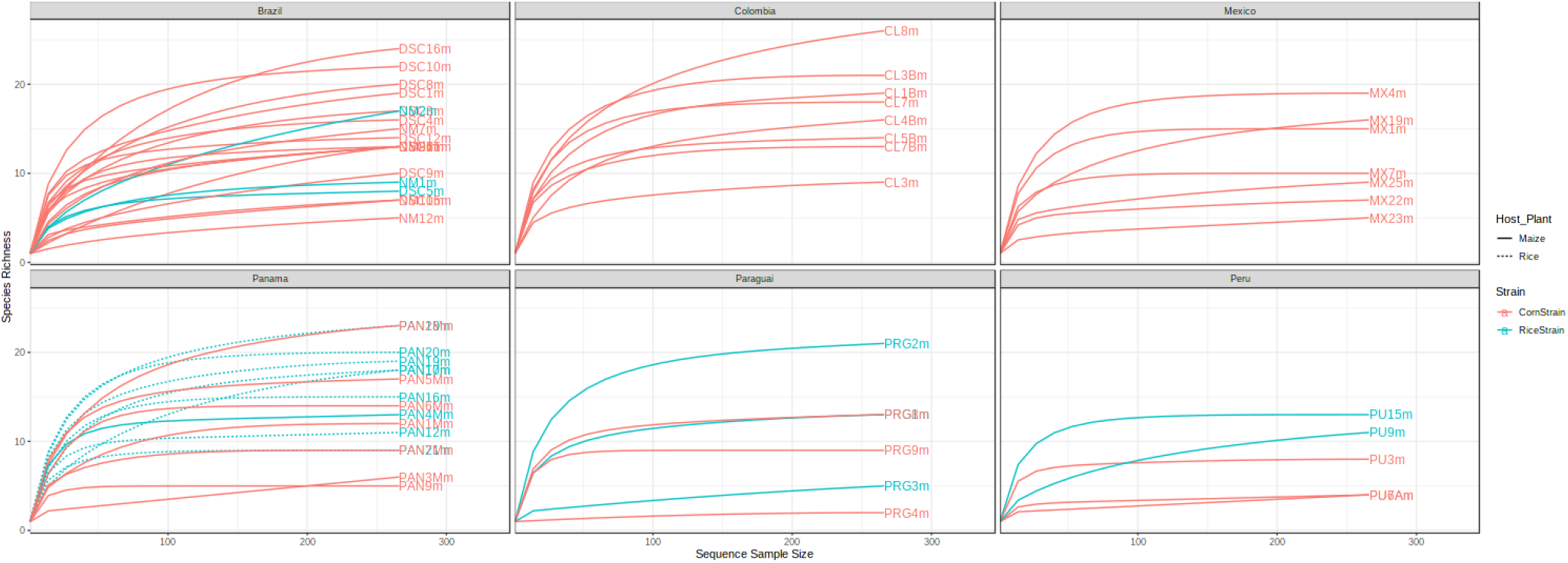
Rarefaction curves showing the relationship between number of ASVs and number of sequences. The rarefaction curve for the midgut of *Spodoptera frugiperda* strains (RS= red and CS=blue) fed on and maize collected in different countries.

